# Comparative genomic and metagenomic investigations of the *Corynebacterium tuberculostearicum* species complex reveals potential mechanisms underlying associations to skin health and disease

**DOI:** 10.1101/2022.08.31.506047

**Authors:** Rauf Salamzade, Mary Hannah Swaney, Lindsay R. Kalan

## Abstract

*Corynebacterium* are a diverse genus and dominant member of the human skin microbiome. Recently, we reported that the most prevalent *Corynebacterium* species found on skin – including *Corynebacterium tuberculostearicum* and *Corynebacterium kefirresidentii* – comprise a narrow species complex despite the diversity of the genus. Here, we apply high-resolution phylogenomics and comparative genomics to describe the structure of the *C. tuberculostearicum* species complex. We find this species complex is missing a fatty acid biosynthesis gene family which is often found in multi-copy in approximately 99% of other *Corynebacterium* species. Conversely, this species complex is enriched for multiple genetic traits, including a gene encoding for a collagen-like peptide. Further, through metagenomic investigations, we find that one species within the complex, *C. kefirresidentii*, increases in relative abundance during atopic dermatitis flares and show that most members of this species possess a colocalized set of putative virulence genes.

## INTRODUCTION

Several species of *Corynebacterium* have been isolated from human skin, often associated with diseases such as diphtheria and regarded as opportunistic pathogens^1,2^. Only recently, with the application of metagenomics to comprehensively survey the skin microbiome, have we begun to develop a better understanding of this understudied genus and assess mechanisms underlying their basis as core constituents of skin microbiomes^3–5^.

Regarded as a lipophilic and disease-associated species, *C. tuberculostearicum* was first named and associated with cases of diphtheria in 1984^1^, later found to be associated with rhinosinusitis^6^, and more recently shown to elicit specific inflammatory signaling pathways of skin cells^5^. While it was noted that *C. tuberculostearicum* could be closely related to other *Corynebacterium* species when it was first validated as a species in 2004^7^, this analysis was based on 16S sequences which provide limited resolution for such investigations. Later work suggested that misclassification of *C. tuberculostearicum* is likely prevalent by clinical biochemical assays and that the species is potentially underreported^8^.

We recently found that the three most prevalent species of *Corynebacterium* in healthy skin microbiomes from a metagenomic survey in our lab^9^, currently classified as *Corynebacterium tuberculostearicum, Corynebacterium kefirresidentii*, and *Corynebacterium aurimucosum* type E in GTDB^10^, belong to a narrow clade, which includes at least one additional species^11^. Because the average nucleotide identity (ANI) between genomes in this clade is relatively high and the lowest value observed was at 88%, we refer to the clade as the *C. tuberculostearicum* species complex^10,11^. While *C. aurimucosum* and *C. tuberculostearicum* have been validated as species^7,12^, reassessment of genomes classified as these species in NCBI’s GenBank database by GTDB suggests that some are mislabelled^10^. *C. kefirresidentii* was only recently proposed as a new species and has not yet been validated by the International Committee on Systematics of Prokaryotes^13^.

Isolates of *C. kefirresidentii* sampled from healthy human skin are lipophilic similar to *C. tuberculostearicum* and also grow better in the presence of compounds commonly found in sweat^9^. This finding is in accordance with our earlier observation that members of the *C. tuberculostearicum* species complex are often found at body sites regarded as moist^11^. In this study, we apply comparative genomics and metagenomics analytics to develop a better understanding of the genetic factors underlying the prevalence of the *C. tuberculostearicum* species complex on healthy skin and determine they lack key metabolic enzymes likely rendering them lipid-dependent. In addition, we assess the biogeographical distribution of individual species within the complex across different skin body sites and whether they associate with atopic dermatitis flares.

## RESULTS

### A high-resolution phylogeny of the *C. tuberculostearicum* species complex

A set of 26 genomes belonging to the *C. tuberculostearicum* species complex were used for phylogenomic investigation of the complex (Table S1). This set of genomes includes 22 genomes we had previously described as part of the complex^11^. In addition, we added three recently reported metagenome-assembled genomes (MAGs) from skin predicted to correspond to novel *Corynebacterium* species^14^ and the genome of a *C. kefirresidentii* we recently isolated from human skin^9^. All 26 genomes, including a total of 8 MAGs, were regarded as high completion and showed low rates of contamination by CheckM analysis^15^ (Table S1).

A set of 1,250 single-copy-core (SCC) orthologs was identified amongst the 26 genomes belonging to the species complex and used to construct a high-resolution phylogeny following filtering of sites predicted to be affected by recent or ancestral recombination^16,17^ (Figure 1). This phylogeny was found to be highly concordant with species classifications for the genomes by GTDB-tk^18^ run with GTDB release 207^10^, but not their taxonomic designations on GenBank (Table S1).

**Figure 1:**
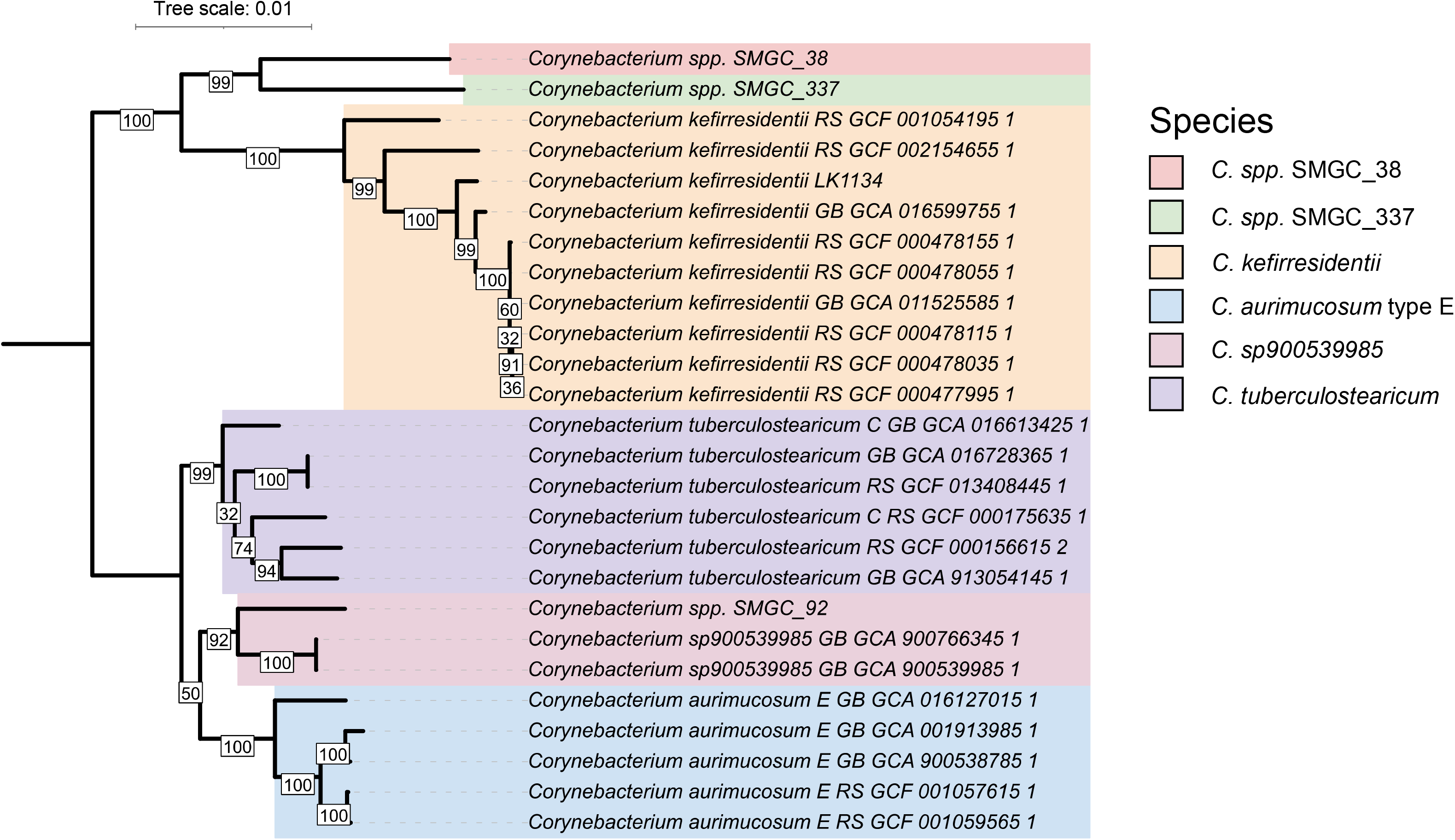
A high-resolution phylogenomic view of the *C. tuberculostearicum* species complex. A maximum-likelihood phylogeny of the *C. tuberculostearicum* species complex, constructed from 15,936 sites predicted to not be impacted by recombination and not exhibit ambiguity in any of the 26 genomes, is shown after midpoint rooting. Bootstrap values are indicated at inner-nodes.

ANI between genomes belonging to different species ranged from a lower limit of ~88%^11^ to nearly 95% (Figure S1). This suggests recent speciation occurring within the species complex, in particular within the subclade containing the species *Corynebacterium* sp900539985, *C. aurimucosum* type E, and *C. tuberculostearicum*, where genomes from different species can exhibit ANI greater than 94% to each other, which is slightly under the 95% ANI threshold commonly used to delineate species^19,20^. In addition, the ANI between genomes *C. spp.* SMGC_38 and *C. spp.* SMGC_337 was estimated as being between 93% and 95%, suggesting recent speciation between them. These two newly reported species^14^, exhibited greater than 90% ANI to *C. kefirresidentii* genomes (Figure S1). The MAG proposed to correspond to a novel *Corynebacterium* species^14^, *Corynebacterium spp.* SMGC_92, was found to exhibit high ANI to an established but currently unnamed species in GTDB, *C.* sp900539985, and designated as belonging to this species. Due to close phylogenetic placement and ANI estimates exceeding 95%, we further propose that GTDB species *C. tuberculostearicum* and *C. tuberculostearicum* type C should simply be regarded as one species (*C. tuberculostearicum).*

Overall, the phylogeny constructed using the core-genome with sites predicted to be affected by recombination removed, was well supported as indicated by high bootstrap values at ancestral nodes to species and sets of species (Figure 1). However, we were unable to confidently resolve whether *C. sp900539985* are more related to *C. tuberculostearicum*, as ANI analysis suggests, or *C. aurimucosum* (Figure 1, S1).

### Unique genes and the biogeographic distribution of the *C. tuberculostearicum* species complex

To address potential biases predicting prevalence of the *C. tuberculostearicum* species complex associated with the metagenomic survey performed in our laboratory^4^, we used StrainGE^21^ to profile the presence of representative *Corynebacterium* genomes amongst published metagenomic datasets profiling the healthy skin microbiome^3^ including a study profiling the skin microbiome in atopic dermatitis, a common pediatric allergic disease^22^. In concordance with our previous findings^11^, we observed that all three species from the complex represented in the StrainGE database are highly prevalent in these additional metagenomic datasets relative to other *Corynebacterium* species (Figure 2A; Table S2). We further observed that *C. tuberculostearicum* and *C. kefirresidentii* were differentially abundant at distinct body sites. While *C. tuberculostearicum* were prevalent and found in high abundances at foot associated body sites, in particular the toe web space, *C. kefirresidentii* were more prevalent in nasal and surrounding alar crease sites (Figure 2B).

**Figure 2:**
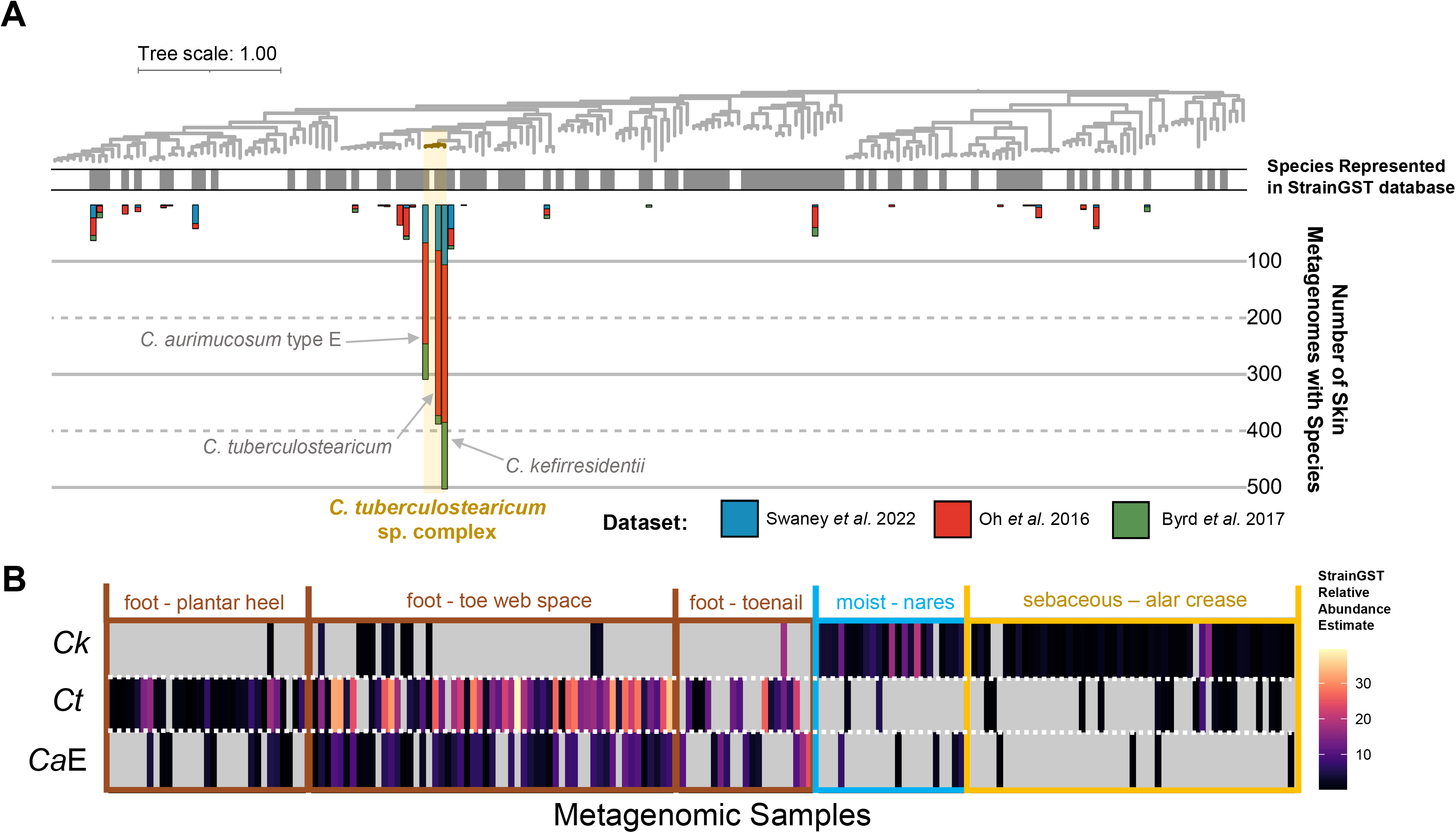
The *C. tuberculostearicum* species complex across skin metagenomic datasets and body sites. **A**) A phylogeny of the 187 *Corynebacterium* is shown with the *C. tuberculostearicum* species complex highlighted in gold. A single track is shown for species which were represented in the StrainGST database which we used to detect strains/species within metagenomes. Bars correspond to the number of skin-related metagenomic samples which were found to feature a species, with colors representing the study the metagenomic. **B**) The estimated relative abundance of *C. kefirresidentii (Ck), C. tuberculostearicum (Ct)*, and *C. aurimucosum* type E (*CaE*) is shown across Oh *et al.* 2016 and Swaney *et al.* 2022 metagenomes from select body sites in adults. Metagenomes without any of the three species detected are not shown.

To understand the basis of the enrichment of the *C. tuberculostearicum* species complex, we performed comparative genomics to determine if particular homolog groups were found more commonly or less commonly than expected within the complex relative to the rest of the genus (Table S3). A representative set of 187 *Corynebacterium* genomes was selected based on dereplication^23^ of those designated as belonging to the genus in GTDB at 95% ANI, to approximate selection of a single representative genome per species.

We identified only two homolog groups which were statistically significant for being absent in the species complex but highly conserved throughout the genus. One of these was annotated as a short-chain dehydrogenase reductase and was absent in all four representative genomes of the *C. tuberculostearicum* species complex but found at a median copy count of three in 98.9% other *Corynebacterium.* Searching for highly conserved domains on the consensus sequence of the homolog group revealed a more precise annotation of 3-oxoacyl-ACP reductase FabG^24^, thus suggesting a role in fatty acid biosynthesis^25^.

A larger set of proteins were identified as being statistically enriched in the *C. tuberculostearicum* species complex in relation to other *Corynebacterium.* We filtered for 26 homolog groups which were found to be nearly core to the species complex (found in >95% of the 26 *C. tuberculostearicum* genomes). While the majority of these homolog groups lacked reliable functional annotations, we identified three to putatively correspond to acetyltransferases and an additional three to be factors in cellular regulation or translation. The largest homolog group, found in only 3.8% of *Corynebacterium* outside of the species complex, was predicted to encode a collagen-like peptide. Collagen-like proteins have previously been suggested to function as adhesins of pathogens to host cells^26–29^ and could correspond to an analogous mechanism in the *C. tuberculostearicum* species complex for attaching to skin cells.

### Species specific partitioning into distinct moist body sites and association with atopic dermatitis

Because prior taxonomic classifications of genomes were incongruent with their phylogenomic placement, it was previously impossible to accurately associate individual species within the *C. tuberculostearicum* species complex with skin-related diseases. With our resolved view of the species complex, we thus reinvestigated the metagenomic dataset from Byrd *et al.* to assess the relative abundance of *C. tuberculostearicum, C. kefirresidentii*, and type E *C. aurimucosum* at different stages of atopic dermatitis for a pediatric cohort. We found that *C. kefirresidentii* was more frequently detected during flares, in comparison to baseline (*p*=1.5e-4) and healthy control microbiomes (*p*=4.0e-4). Furthermore, in certain metagenomes sampled during a flare, *C. kefirresidentii* exhibited a substantial increase in relative abundance (Figure 3A; S2).

**Figure 3:**
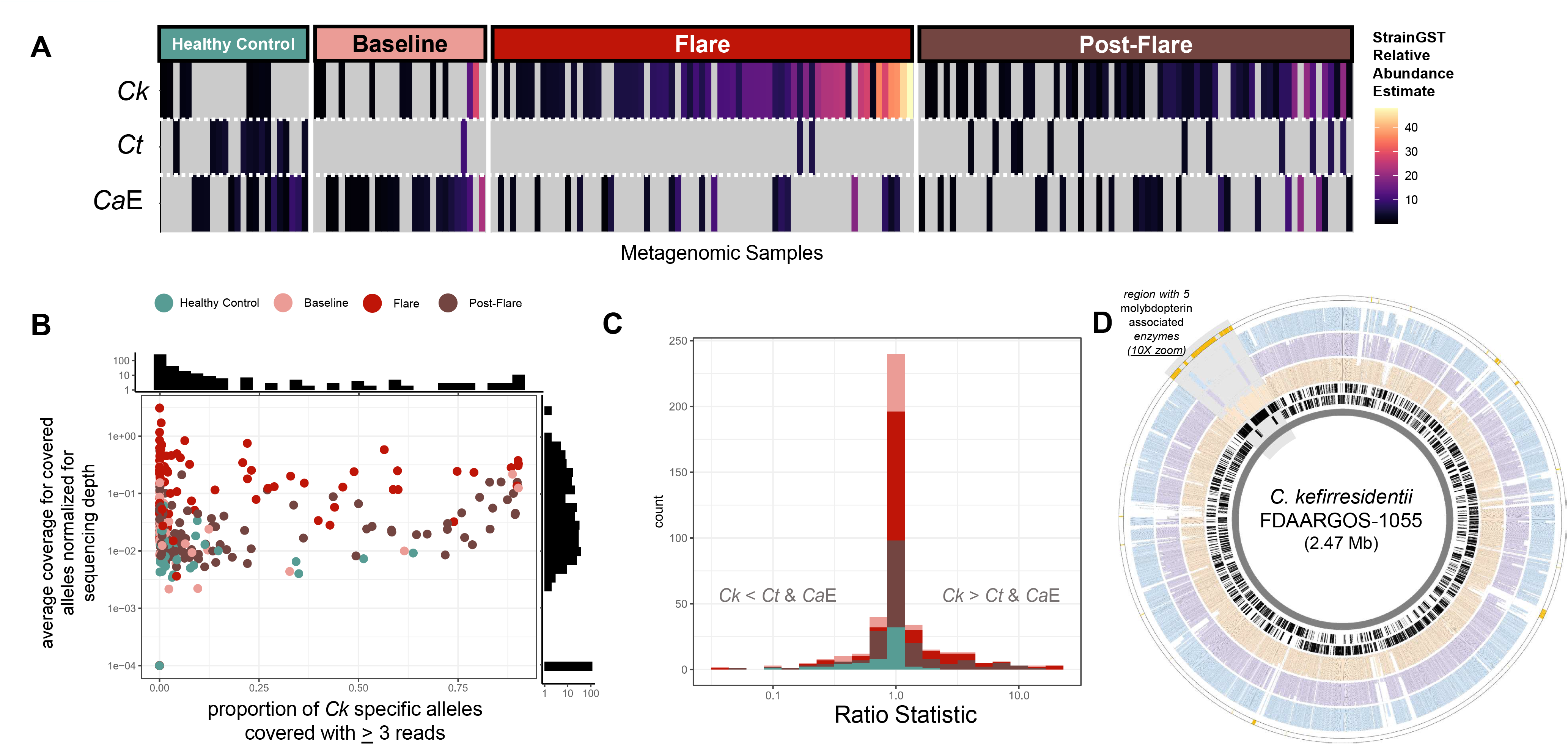
Increase in *C. kefirresidentii* relative abundance in metagenomes sampled during atopic dermatitis flares. **A**) The estimated relative abundance of *Corynebacterium* species in the species complex is shown across all metagenomes from Byrd *et al.* 2017 with at least one species from the complex detected. **B**) Average coverage of *C. kefirresidentii* specific alleles along 367 single-copy-orthologs of the *C. tuberculostearicum* species is shown. Only sites/alleles which were covered by at least three reads are regarded. **C**) The ratio of the total average coverage for *C. kefirresidentii* specific alleles (inclusive of uncovered alleles) to the sum of total average coverage for *C. tuberculostearicum* specific alleles and *C. aurimucosum* type E specific alleles is shown. A pseudo-count of 1 was added to both the numerator and denominator. **D)** Genomic location of homolog groups identified as enriched in *C. kefirresidentii* genomes relative to *C. tuberculostearicum* and *C. aurimucosum* type E genomes along the chromosome of *C. kefirresidentii* FDAARGOS_1055 is shown as the outermost track in gold. The most inner two rings in black show the location of coding genes on the reverse and forward strands. The following rings (from inner to outer) show the conservation of homologs of the gene in *C. kefirresidentii* (orange), *C. tuberculostearicum* (purple), and *C. aurimucosum* type E (blue) genomes. The co-located set of genes found to be conserved in 80% of *C. kefirresidentii* featuring five molybdopterin associated enzymes is highlighted in grey and zoomed in at 10X.

To further affirm the association of *C. kefirresidentii* with atopic dermatitis flares, we also applied an alignment-based approach in which we aligned metagenomic reads to the single-copy core genes of the complex and measured coverage of species specific alleles. *C. kefirresidentii* was more evidently present at higher abundances primarily in metagenomes from flares and post-flare samplings (Figure 3B), which also tended to have the lowest viable sequencing data after filtering metagenomes for quality, adapter sequences, and human contamination (Figure S3). We further developed a ratio statistic to assess the abundance of *C. kefirresidentii* to *C. tuberculostearicum* and type E *C. aurimucosum* individually within each metagenome. *C. kefirresidentii* were more abundant than type E *C. aurimucosum* and *C. tuberculostearicum* combined in 52.7% of metagenomes survey from flares, which was significantly more than for baseline (25.7%; *p*=2.3e-4) and healthy control (21.1%; *p*=3.8e-5) metagenomes (Figure 3C).

To understand what potential genetic factors might be contributing to the unique association of *C. kefirresidentii* with atopic dermatitis flares, we performed comparative genomics to identify traits which were lost or gained in their genomes relation to the closely related *C. tubercuostearicum* and type E *C. aurimucosum* species. We found a total of 97 homolog groups were either enriched or depleted in *C. kefirresidentii* relative to *C. tubercuostearicum* and type E *C. aurimucosum* (Table S4). *C. tubercuostearicum* and type E *C. aurimucosum* feature a core thermonuclease that is homologous to the virulence factor *nu^c30^* from *Staphylococcus aureus*, but missing in all *C. kefirresidentii.* In contrast, two distinct colocated sets of genes, one predicted to encode for thiamine biosynthesis and the other encoding molybdopterin-binding enzymes, were enriched in *C. kefirresidenttii* but missing or significantly lacking in *C. tubercuostearicum* and type E *C. aurimucosum* genomes. Of note, molybdenum enzymes have recently been discussed as important virulence factors for infection in a variety of diverse pathogens^31,32^.

## DISCUSSION

The *C. tuberculostearicum* species complex features the three major species of *Corynebacterium* found on human skin^11^. Despite this, fewer than 30 representative genomes across all species belonging to the complex are present in the most recent GTDB release and eight of the 26 genomes (30.8%) used in this study were MAGs. The lack of genomes for the species complex, despite its prevalence on healthy skin, might be in part due to difficulties with culturing. For instance, we recently found that a single *C. kefirresidentii* isolate grows best when high concentrations of compounds found in both sebum and sweat are available^9^.

Here we determine that the *C. tuberculostearicum* species complex appears to be missing a fatty acid biosynthesis gene family that is often found in multi-copy in other *Corynebacterium*, potentially explaining their lipophilic nature^7,8^. However, other *Corynebacterium* species outside of this species complex have also been characterized as lipophilic^9,33^, suggesting convergent evolution with different genetic factors underlying the shared trait in diverged species across the genus. Comparative genomics further highlighted several genes encoding proteins that are specific to the species complex, though the majority of these lack any functional annotation and future molecular characterization will be essential to further our functional understanding of them. Interestingly, one of these genes, core to the species yet present in less than 4% of other *Corynebacterium*, encodes for a collagen-like peptide and could be involved in adherence to skin.

We further identify an increase in relative abundance of *C. kefirresidentii* in metagenomes sampled during atopic dermatitis flares. Comparative genomics between *C. kefirresidentii* with *C. tuberculostearicum* and *C. aurimucosum* type E genomes found that multiple genes differentiate *C. kefirresidentii* from its two neighboring species. Most striking are a set of five colocated molybdopterin-associated enzymes, which are commonly associated with virulence^31,32^.

Two ATCC isolates (ATCC-35693 / 2628 LB and ATCC-35694 / FPSA)^34^, designated as *C. tuberculostearicum* and originating from the original study first describing the species taken from leprosy patients^1^, have more recently been used in studies to associate the species to rhinosinusitis^6^ and elicitation of a specific host inflammatory signaling pathway^5^. To our knowledge, these isolates lack genomes however and might currently be misclassified since other species of *Corynebacterium* have been reported to similarly produce tuberculostearic acid, the trait initially used to differentiate the species from others in the genus^7,12,35^. Identifying genetic markers to distinguish species within the *C. tuberculostearicum* species complex and continued efforts in whole genome and metagenomic sequencing will thus be critical to correctly associate the contributions of particular species within the complex to diseases.

## ACKNOWLEDGMENTS

This work was supported by grants from the National Institutes of Health (NIAID U19AI142720 and NIGMS R35GM137828 [L.R.K]). The content is solely the responsibility of the authors and does not necessarily represent the official views of the National Institutes of Health. The authors gratefully acknowledge J.Z. Alex Cheong, Dr. Nicole Lane Starr, Lucas van Dijk, Dr. Abigail Manson, and Dr. Ashlee Earl for helpful discussions.

## METHODS

### Genome selection and taxonomic reclassification

We had previously identified a set of 22 genomes through an interactive process in which we first identified the clade using phylogenomics as consisting of the five GTDB defined species: *C. tuberculostearicum, C. tuberculostearicum_C, C. aurimucosum_E*, and *C. kefirresidentii* (using release R202)^11^. Five additional genomes belonging to the clade were identified in NCBI which were missing in GTDB release R202 and validated to belong to one of the five species through GTDB-tk classification^18^. In our earlier study, we also found that three recently published MAGs from skin microbiomes, proposed novel species of *Corynebacterium* (based on GTDB release 88)^14^, had >88% ANI to one of the prior defined 22 genomes. We thus included these three genomes, in addition to the genome for a *C. kefirresidentii* isolate, LK1134, we had isolated from healthy skin and profiled the growth dynamics off across different concentrations of artificially synthesized sebum and sweat^9^.

Systematic reclassification on the total set of 26 genomes proposed to belong to the *C. tuberculostearicum* species complex was performed using GTDB-tk2 with the latest GTDB release R207. All genomes were assigned to one of the five previously mentioned GTDB species, except for the MAGs *C. spp.* SMGC_38 and *C. spp.* SMGC_337^14^, which appear to still correspond to novel species of close relation to *C. kefirresidentii.* Because MAGs were included, the contamination and completeness of genomes was assessed using CheckM (v1.2.1) (Table S1).

For comparative genomics of the *C. tuberculostearicum* species complex against other species from the *Corynebacterium* genus, all 1,296 genomes in the genus from GTDB R207 were downloaded^10^. Genome dereplication was performed using dRep^23^ with a secondary ANI filter of 95% using fastANI^19^. This retained a set of 187 distinct genomes, including four representatives of the *C. tuberculostearicum* species complex.

### Homolog group inference and functional annotation

Gene calling was performed on genomes using prodigal^36^ and OrthoFinder (v2.5.4^)16^ was subsequently used to perform homolog group detection independently amongst the set of 26 *C. tuberculostearicum* genomes and the genus-wide set of 187 representative *Corynebacterium* genomes. For both analyses, the course set of initial orthogroups following MCL were used instead of phylogenetic hierarchical orthogroups.

Functional annotation was only performed for homolog groups which comparative genomics investigations suggested were significantly more prevalent or lacking within certain phylogenetic clades. Annotation was primarily performed using individual protein sequences from such homolog groups via the EggNOG mapper webserver run with default settings^37^. Consolidation of annotations was performed across proteins belonging to each homolog group. For homolog groups predicted to be enriched or depleted in the *C. tuberculostearicum* species complex relative to other *Corynebacterium*, we further attempted annotation by generating consensus amino acid sequences for each homolog group and using NCBI’s conserved domain search^24^ as well as Phyre2 to assess structural similarity to structurally characterized proteins^38^.

### Phylogenomics of the *C. tuberculostearicum* species complex and the representative selection of *Corynebacterium*

We identified 1,250 homolog groups as single-copy orthologs between the 26 *C. tuberculostearicum* species complex genomes. Protein alignments were constructed using MUSCLE (v5)^39^ and converted to codon alignments using PAL2NAL^40^. RAxML (v8.2.12) was then used to construct an initial maximum likelihood phylogeny on a concatenation of codon alignments (1,239,366 bp) with the GTRCAT model and 1,000 bootstraps. Core codon alignments were projected onto and ordered according to the reference genome *C. kefirresidentii* FDAARGOS_1055 and, together with the RAxML phylogeny, used as input for ClonalFrameML (v1.12)^17^ to infer sites affected by recombination by first using a simple model, with parameter emsim set to 100 and kappa (the relative rate of transitions to transversions) set to 5.09664. For reference projection, gaps between single-copy orthologs were set to be the length of distances in between corresponding genes in the reference genome and codon alignments were reverse complemented if the gene was predicted to occur on the antistrand. Non-core sites were specified as the gaps placed in between single copy core orthologs so as to appropriately model the strength of linkage between them. Results from running ClonalFrameML with a simple model were used to initialize values for R/theta, 1/delta, and nu (set to 0.126997, 0.0027686, and 0.0583416, respectively) and run ClonalFrameML with a per-branch model with the embranch_dispersion parameter set 0.1, to allow for greater dispersion in parameters amongst branches. The initial alignment was filtered for sites predicted to be affected by recombination and subsequently subset for 15,936 sites which lacked ambiguity for any of the 26 genomes. This alignment was finally used to construct a high-resolution, maximum likelihood phylogeny of the species complex using RAxML with the same modeling strategy as applied initially.

For phylogeny construction of the full genus, we used GToTree^41^ (v1.6.36) with the set of single-copy gene sets specific to Actinomycetota (formerly Actinobaceria) and FastTree2 phylogeny inference^42^.

### Statistical testing for enrichment or depletion of homolog groups in clades for comparative genomics analyses

Assessment of enrichment or depletion of homolog groups for two sets of genomes was carried out using a two-sided permutation test based on mean copy-count differences with 100,000 resamples. This allowed us to detect homolog groups which were not merely differentially present, but also those which had greater copy-counts in one set of genomes relative to the other. A generalizable program for this testing is provided in the Github repository: https://github.com/Kalan-Lab/Salamzade_etal_CtuberCompGen. Multiple testing correction was performed for both such comparative genomics analyses using the Benjamini-Hochberg procedure for P-value adjustment to control the false discovery rate. For homolog groups found to be enriched in the four *C. tuberculostearicum* species complex representative genomes relative to the complementary set of 183 *Corynebacterium* genomes, we further required that they map to homolog groups deemed to be >95% near-core amongst the 26 *C. tuberculostearicum* species complex genomes (there was a total of 1,871 near-core homolog groups).

### Metagenomic detection of *Corynebacterium* strains and species

Three independent sets of metagenomes profiling the microbiome of healthy skin and skin affected by atopic-dermitis were processed using a pipeline previously described^4^. Briefly, metagenomes were processed for quality and human contamination using fastp^43^ and KneadData (https://github.com/biobakery/kneaddata). Mock and negative control metagenomes from the Swaney *et al.* 2022 study were excluded^4^ as were 22 metagenomes from the Oh *et al.* 2016 dataset marked as “human depletion” or “whole genome amplification” under “Notes (excluded for comparisons)” in their spreadsheet describing all samples^3^. Deeply sequenced metagenomes from Oh *et al.* 2016 were randomly downsampled to 25 million read pairs using seqtk (https://github.com/lh3/seqtk). In addition, as described previously^4^, five metagenomes from the Byrd *et al*. 2017 dataset^22^ (3 flare and 2 post-flare) were excluded due to insufficient sequencing data following processing. All symmetric metagenomes (e.g. metagenomes corresponding to sampling the left and right sides of a body site) from the Oh *et al.* 2016 and Byrd *et al.* 2017 metagenomes were individually considered.

Detection of strains or representative genomes was performed using StrainGST^21^ with a custom database constructed from complete *Corynebacterium* genomes from NCBI, as described previously^11^. Briefly, this database featured 164 distinct representative genomes following dereplication using programs for StrainGST (0.1+191.g7fcfbcd.dirty) database creation^21^. These genomes were run through GTDB-tk with database release R207^10,18^ to generate consistent taxonomic classifications. One genome regarded as *Corynebacterium* in RefSeq^44^ at the time of database creation was classified as not part of the genus by GTDB-tk and not observed in any of the metagenomic datasets. Another genome was classified as a novel *Corynebacterium spp.* not yet given an identifier in GTDB R207. We counted the number of distinct species detected in metagenomes from the three datasets and illustrated them as a multi-bar graph on the phylogeny of 187 *Corynebacterium* by species name. Because *C. tuberculostearicum* and *C. tuberculostearicum* type C classified genomes were found in this study to phylogenetically group as one, we considered them as a single species and mapping of detection counts for the *C. tuberculostearicum* representative genome in StrainGST was allowed to the *C. tuberculostearicum* type C representative genome in the set of 187 genomes used for phylogeny construction and comparative genomics. The species *C. sp900539985* from the *C. tuberculostearicum* species complex, which was represented in the phylogeny of 187 genomes, was not represented in the StrainGST database as it lacks a complete genome currently. Further, there was a single representative strain / genome for each of the three species from the complex represented in the StrainGST database: *C. tuberculostearicum*, *C. aurimucosum_E*, and *C. kefirresidentii.* The relative abundance estimation for these three species was thus computed based on the relative abundance computed for the corresponding representative strains / genomes by StrainGST *(rapct)* after updating to version 1.3.3, to incorporate updates to how relative abundance was being computed relative to earlier versions.

#### Confirmation of StrainGST results using an independent approach

StrainGST is a k-mer based approach to assess strain presence in metagenomes and to complement it we used an independent and complementary alignment based approach to gauge whether *C. kefirresidentii* were enriched during atopic dermatitis flares in the Byrd *et al.* 2017 metagenomes. For this approach, we aligned metagenomic reads individually to 4,514 representative genes from 367 homolog groups which comparative genomics had revealed were single-copy orthologs and featured at least 20 sites where *C. kefirresidentii* had a core allele that was not observed in the complementary set of genomes in the *C. tuberculostearicum* species complex. Reads were aligned as unpaired using Bowtie 2^45^ with parameters “--very-sensitive-local --no-unal -a -x”. Reads were aligned as unpaired because gene sequences are short and pairs can thus align poorly. Additionally, all alignments for each read are requested because multiple representative genes were considered per homolog group. We then processed alignments to each representative gene where coverage at 1X was not observed for 90% of the sites. Following this preliminary scan for checking that genes were loosely covered, we assessed alignments to the gene met either of the following two criteria: (i) >99% identity with length >60 bp or (2) >95% identity with length >100 bp for the core alignment region (excluding flanking regions to account for alignments hanging off representative gene edges). We also allowed a maximum of 5 indelic sites within the core alignment region for the alignment to be considered. Alignments which met these criteria were then further processed and retained if they had the best mapping score observed for a particular read (in the case of ties, all alignments with the best Bowtie 2 mapping score were retained). Finally, allele counts were tallied for each metagenome at each site in homolog group codon alignments based on mapping of the query base to the reference / representative gene position in the codon alignments. Only positions in reads which had greater than 30 base quality were considered.

Results from determining the presence of alleles at sites along homolog group codon alignments for each metagenome were processed to identify the proportion of *C. kefirresidentii* specific alleles covered by at least three reads. The mean coverage of these well-covered *C. kefirresidentii* specific alleles were computed and normalized for sequencing depth through dividing by a metagenome specific scalar value (Figure S3). This scalar value corresponded to the ratio of the number of bases the metagenome featured in relation to the metagenome with the fewest number of bases in the final forward read files following processing.

We additionally identified *C. tuberculostearicum* and type E *C. aurimucosum* specific alleles along the same set of 367 core homolog groups and quantified the mean coverage of species specific alleles within each metagenome (inclusive of coverage at alleles which were uncovered). This allowed us to compute a ratio statistic measuring the abundance of *C. kefirresidentii* to *C. tuberculostearicum* and type E. *C. aureimucosum* with a pseudocount of 1 added to both the numerator and denominator to avoid dividing by zero. A value greater than 1 for this statistic thus indicates that *C. kefirresidentii* are more prevalent than *C. tuberculostearicum* and type E. *C. aureimucosum* within an individual metagenome; whereas, a value less than 1 supports the opposite. Our code for metagenomic alignment and calculation of the ratio statistic is provided in the Github repository: https://github.com/Kalan-Lab/Salamzade_etal_CtuberCompGen.

#### Statistical testing

To test whether StrainGST-based detection of *C. kefirresidentii* was significantly more prevalent within metagenomes in atopic dermatitis flare metagenomes relative to baseline and healthy control metagenomes, we used the two-sided Fisher’s exact test. The two-sided Fisher’s exact test was also used to statistically assess whether the ratio statistic was larger than 1 in metagenomes sampled during atopic dermatitis flares relative to baseline and healthy control metagenomes.

### Visualizations

iTol was used for illustrating phylogenetic trees and associated features^46^. The R library ggplot2^47^ was used to visualize heatmaps, scatterplots, and bar graphs. Circleator^48^ was used to generate an overview of conservation and showcase genes highlighted by comparative genomics along the chromosome of *C. kefirresidentii* FDAARGOS_1055.

