## Supplementary Figures S1 to S3 for "Comparative genomic and metagenomic investigations of the *Corynebacterium tuberculostearicum* species complex reveals potential mechanisms underlying associations to skin health and disease"

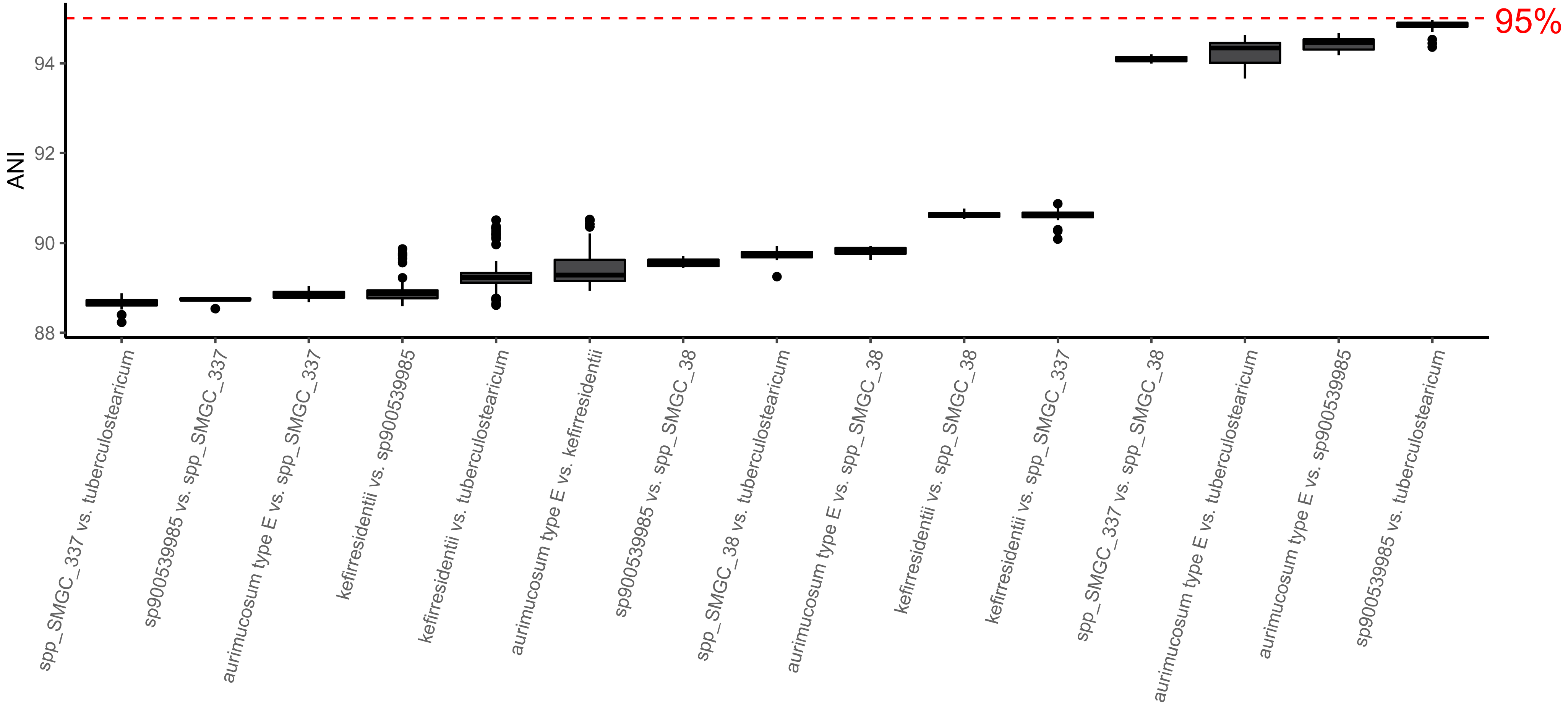


**Figure S1: ANI estimates between species in the *C. tuberculostearicum* species complex.** fastANI was used to estimate ANI between the four species within the *C. tuberculostearicum* species complex and the MAGs SMGC38 and SMGC337, which are predicted to represent closely related novel species not represented in GTDB R207.

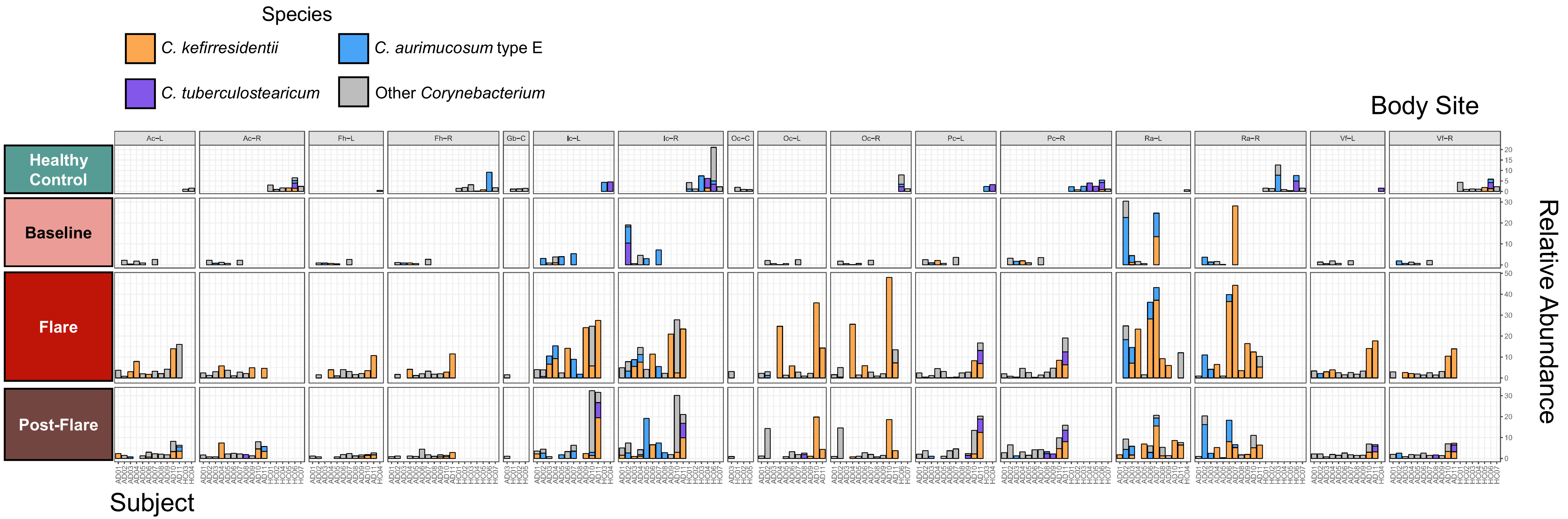


**Figure S2: Increase in *C. kefirresidentii* relative abundance is observed at multiple body sites during flares.** The relative abundance of *C. tuberculostearicum*, *C. aurimucosum* type E, *C. kefirresidentii,* and other *Corynebacterium* represented in the StrainGST database is shown for healthy control, baseline, flare, post-flare metagenomes from Byrd *et al.* 2017. Vertical panels correspond to body-sites (Ac = antecubital crease, Fh = forehead, Ic = inguinal crease, Oc = occiput, Pc = popliteal crease, Ra = retroauricular crease, Vf = volar forearm), with –R and –L corresponding to sampling taken from the right and left sides of the corresponding body site, respectively.

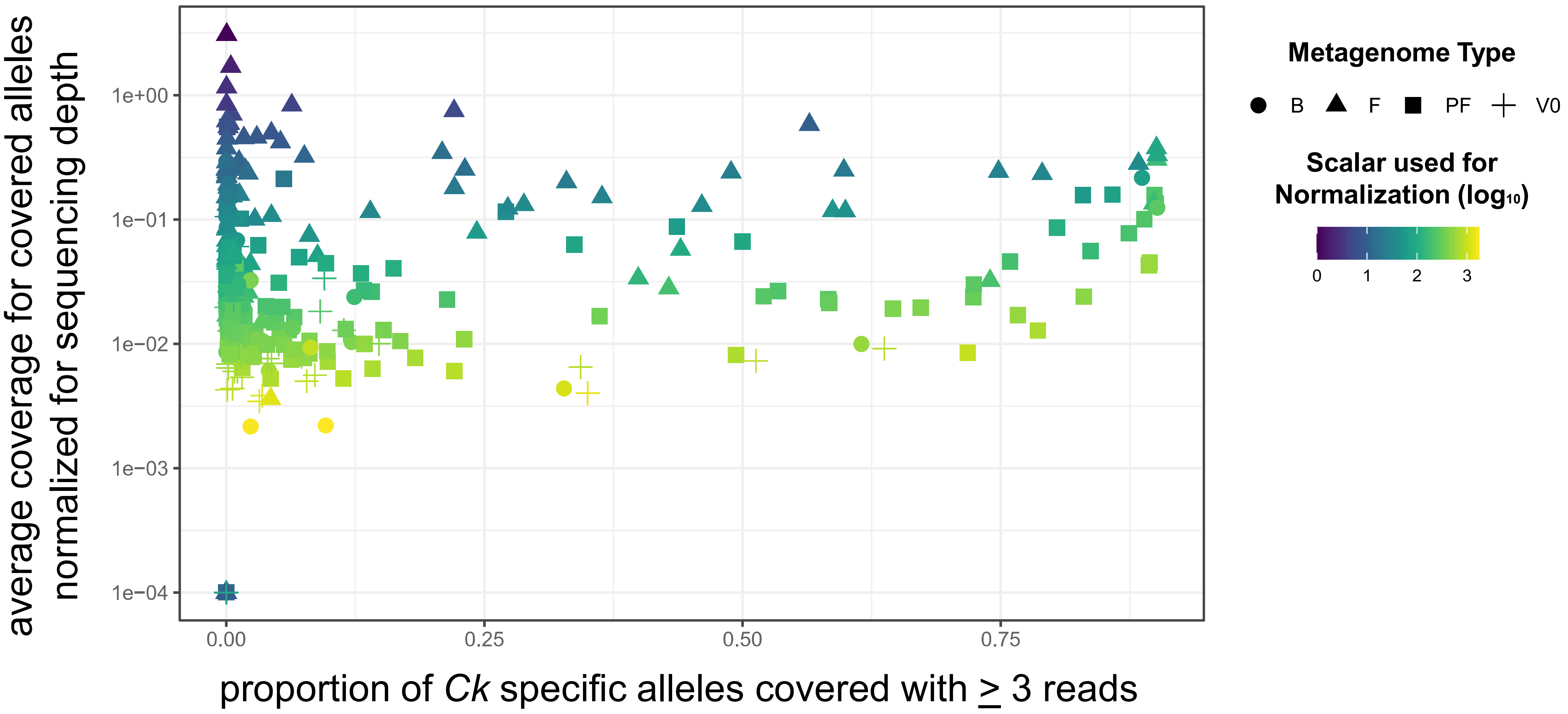


**Figure S3: Flare metagenomes from Byrd *et al.* 2017 have fewer sequencing data after post-processing.** This figure is analogous to Figure 2B, but with shapes corresponding to the four types of metagenomes (B=baseline, F=flare, PF=post-flare, and V0=healthy control) and coloring corresponding to the scalar used for normalization of the y-axis for metagenome sequencing depth post-processing.
